# Local Targeted Memory Reactivation in Human Sleep

**DOI:** 10.1101/539114

**Authors:** Ella Bar, Anat Arzi, Ofer Perl, Ethan Livne, Noam Sobel, Yadin Dudai, Yuval Nir

**Affiliations:** Department of Neurobiology, Weizmann Institute of Science, Rehovot, 76100, Israel; Department of Psychology, University of Cambridge, Cambridge, UK; Department of Physiology & Pharmacology, Sackler School of Medicine, Tel Aviv University, Israel; Sagol School of Neuroscience, Tel Aviv University, Israel; Functional Neurophysiology and Sleep Research Lab, Tel-Aviv Sourasky Medical Center, Tel Aviv 64239, Israel

**Keywords:** TMR, olfactory, odor, NREM, slow waves, spindles, slow oscillations, sleep oscillations

## Abstract

Memory consolidation can be promoted via Targeted Memory Reactivation (TMR) that re-presents training cues or context during sleep. Whether TMR acts locally or globally on cortical sleep oscillations remains unknown. Here we exploit the unique functional neuroanatomy of olfaction with its ipsilateral stimulus processing to perform local TMR in one brain hemisphere. Participants learned associations between words and locations in left or right visual fields with contextual odor throughout. During post-learning naps, odors were presented to one nostril throughout NREM sleep. We found improved memory for specific words processed in the cued hemisphere (ipsilateral to stimulated nostril). Unilateral odor cues locally modulated slow wave activity (SWA) such that regional SWA increase in the cued hemisphere negatively correlated with select memories for cued words. Moreover, local TMR improved slow wave-spindle coupling specifically in the cued hemisphere. Thus, TMR in human sleep transcends global action by selectively promoting specific memories associated with local sleep oscillations.

## Introduction

Ample evidence suggests that sleep is critical for memory consolidation, in which new and labile memories encoded in wakefulness are transformed into less-labile representations that become integrated into pre-existing knowledge [1–6]. It has been confirmed that sleep-related benefits in memory performance are not simply due to reduced fatigue or interference or a distinct circadian state [1]. Sleep supports multiple types of subsequent memory performance, including hippocampal-dependent declarative memory and object-place associations, procedural memory, perceptual, and emotional memory (reviewed in [1]).

According to the standard model of systems consolidation, as well as to some of its subsequent variants [2, 3], consolidation of hippocampal-dependent memory occurs in a two-stage process, whereby new memories are initially encoded into the hippocampus during wakefulness, and subsequently consolidated in a process that involves cross-talk with neocortex, mostly during non-rapid eye movement (NREM) sleep [4]. The hippocampal-neocortical dialogue during sleep likely involves coordinated coupling between neocortical slow waves (SWs), thalamo-cortical sleep spindles, and hippocampal sharp-wave ripples (SWRs). SWRs co-occur with reactivation of neuronal ensembles that were selectively engaged in the learning phase [1, 5, 6]. According to this model, cortical slow-wave up-states initiate hippocampal ripples and neuronal reactivations [7]. Hippocampal reactivation triggers a delayed cortical response that facilitates the transformation to gradually augmented neocortical dependence [8]. Sleep spindles prime the recipient cortical circuits for plasticity [9], ensuring optimal reception. Thus, hierarchical nesting of SW-spindle-SWR events may promote the transformation of reactivated hippocampal memories to respective neocortical networks [5].

Recent work suggests that memory consolidation during NREM sleep can be externally modulated by re-presentation of contextual cues (e.g. a specific odor or sound associated with select items during learning) [10–12]. This method, known as “Targeted Memory Reactivation” (TMR), promotes memory consolidation and induces hippocampal activity [10], suggesting that it involves reactivation of the fresh engram or part of it. TMR has been implemented successfully in a variety of declarative and non-declarative memory tasks [10, 13–15]. Although odor and sound both serve as effective contextual stimuli, odor entails some advantages for TMR since it rarely wakes up sleeping participants [16–18]. In addition, odors are powerful cues for memories [19, 20], possibly due to direct projections from the olfactory cortex to the hippocampus that largely bypass the thalamus [21, 22].

However, a major unresolved issue in TMR research is whether selective memory benefits entail local or global effects on cortical sleep oscillations. To date, potential local effects were difficult to reveal given that sensory cueing during sleep also modulates cortical sleep oscillations globally [23–25]. In addition, typical TMR research compares cortical oscillations across different sleep intervals, inevitably introducing variability in sleep activities that masks potential local modulations. To overcome existing limitations in TMR research one would ideally cue specific stimuli that trigger processing in select cortical regions, while simultaneous activity in other areas serve as control. Recent studies established the notion of local sleep oscillations and their relation to memory consolidation [26–34]. Thus, in principle, cortical sleep oscillations may differ across regions that undergo TMR. But, can one modulate human memories in a regional manner?

To perform TMR for specific memories associated with activity in select brain areas, we took advantage of the unique functional neuroanatomy of the olfactory system, where stimulus processing is largely ipsilateral [21, 35]. Although odor information does reach the contralateral hemisphere [36, 37], the first interhemispheric connections in mammals occur at the olfactory cortex. Interhemispheric connections emerge in the anterior olfactory nucleus (AON) and the piriform cortex, and cross to the other hemisphere via anterior commissural pathways [21, 35, 37, 38] but see [39]. Accordingly, olfactory memories can be stored, at least temporally, in one hemisphere, accessible to the other hemisphere only through commissural pathways [40]. Indeed, in newborn rats, memory for learned odor is confined to one hemisphere if a single nostril is stimulated during learning (paring odor with milk reward). The pups show increased preference for the learned odor only if presented to the same nostril that was open at learning. This phenomenon ceases when the pups reach the age of 12 days, simultaneously with maturation of the anterior commissure [40, 41]. Given that during NREM sleep, effective connectivity across cortical regions is reduced [42], we hypothesized that integration across cortical hemispheres during sleep may be jeopardized in a manner akin to an undeveloped commissure. In other words, we predicted that presenting contextual odor cues to a single nostril in sleep would benefit memories and modulate sleep oscillations associated with the ipsilateral hemisphere.

Based on the aforementioned rationale, we developed a novel method to locally reactivate memories in a single hemisphere, using odor stimulation to a single nostril during sleep. This approach allows to assess the behavioral and electrophysiological influences of local cueing on memories related to one hemisphere, while memories and activities related to the other hemisphere serve as control (Fig. 1A). We used a spatial memory task that requires encoding of paired associations between words and specific locations in either the left or the right visual hemifields, associated with processing in the contralateral cerebral hemisphere. The results establish that local TMR in sleep via unilateral olfactory stimulation goes beyond global effects and elicits selective memory enhancement associated with regional differences in cortical sleep oscillations across the cued and uncued hemispheres.

**Figure 1.**
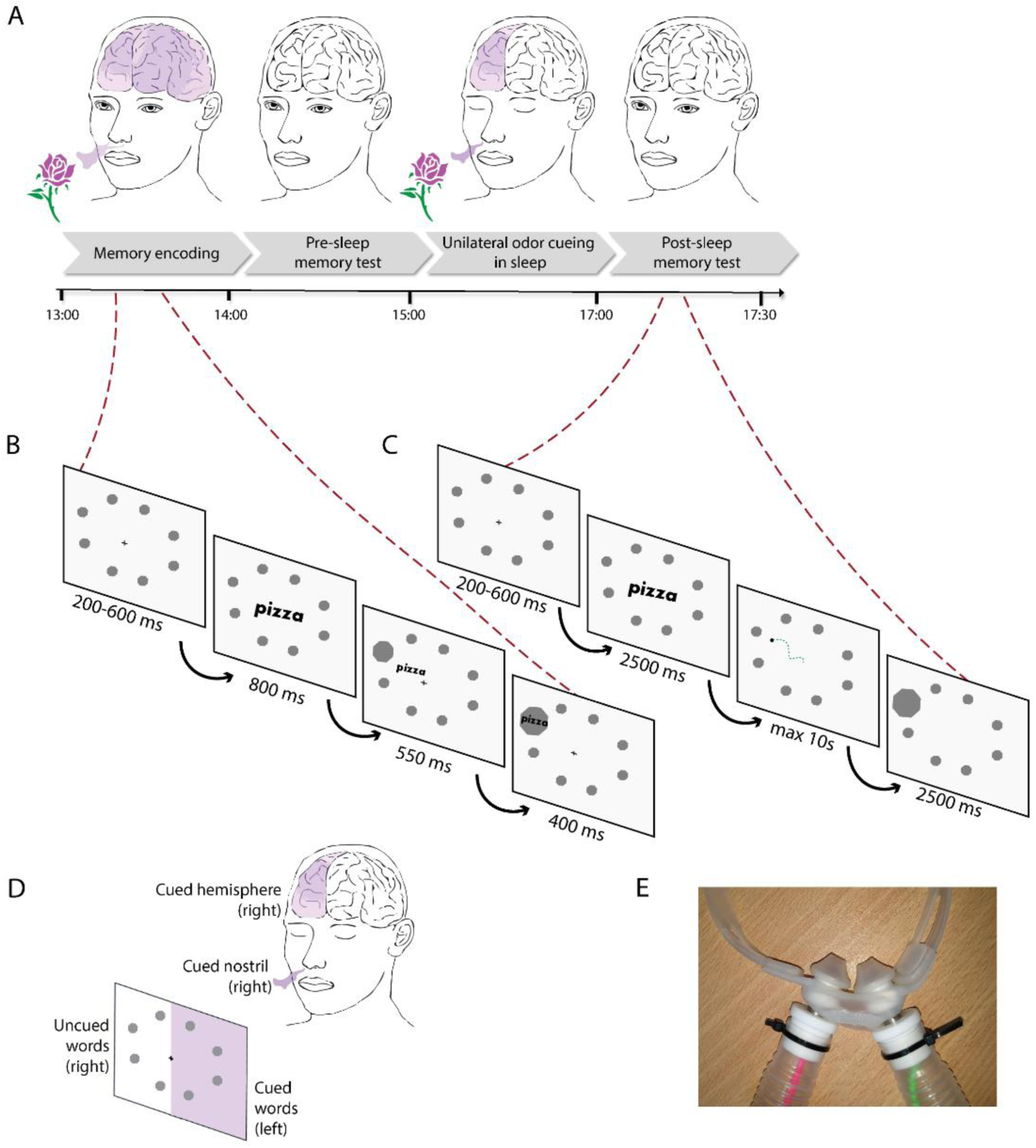
Experimental design. (A) Schematic timeline illustrating the experimental procedure. Participants first learned word-location associations in the presence of a rose odor delivered via mask to both nostrils (“Memory encoding”). Immediately after learning, memory retrieval was tested (“Pre-sleep memory test”). Next, participants took a short (~1-1.5h) nap in the lab. During NREM sleep, odor was delivered unilaterally via special mask to a single nostril (“Unilateral odor cueing in sleep”). Finally, a post-sleep test of memory retrieval was performed (“Post-sleep memory test”). (B) Learning trial timeline. Words appeared sequentially at the center of the screen and moved towards one of eight circular targets. (C) Memory retrieval trial timeline. Words appeared sequentially at the center of the screen, and participants chose the target location associated with each word using the cursor. Green dashed line illustrates an example cursor trajectory. (D) The nasal sleep mask enabling unilateral odor stimulation. (E) Illustration of the terminology: Odor cues delivered during sleep to the cued nostril (in this example, the right nostril; anatomical sides were counterbalanced across participants). The ipsilateral hemisphere is the cued hemisphere (olfactory pathways are mainly ipsilateral). Words associated during learning with target locations in the hemifield contralateral to the cued hemisphere’ (in this example, targets in the left hemifield) are cued words (visual pathways are mainly contralateral).

## Results

We performed local TMR via unilateral olfactory stimulation in sleep to test how it may differentially affect memory for cued items and corresponding regional sleep oscillations. To this end, we presented odors that served as contextual cues during pre-sleep object-location associative learning to a single nostril during sleep (Fig. 1). We use the terms cued/uncued hemisphere throughout to refer to the hemisphere ipsilateral/contralateral to the stimulated nostril during sleep, respectively, regardless of the anatomical cueing side (left/right, counterbalanced across participants). The terms cued/uncued words denote word stimuli presented at the visual hemifield contralateral/ipsilateral to the cued hemisphere, respectively, reflecting our prediction that unilateral cueing will improve memory consolidation for items primarily processed in the hemisphere of the stimulated nostril, presented at the contralateral visual hemifield (Fig. 1D). This terminology is used for description of the procedure but does not imply that processing of odors or individual words is exclusively confined to one hemisphere.

### Local TMR during sleep selectively improves hemisphere-related memories

Memory for word-location associations was evaluated in two sessions: immediately (~5 min) after training (pre-sleep) and ~2.5h later, following a nap (post-sleep). To test for selective TMR effects on memory for cued words, we conducted a two-way repeated measures ANOVA with conditions of time (pre-sleep/post-sleep) and cueing (cued/uncued). We found no main effect of time (F(1,18)=2.85, p=0.1), no main effect of cueing (F(1,18)=0.05, p=0.826), but a significant interaction between time and cueing (F(1,18)=8.58, p=0.009). Post-hoc comparisons revealed that this interaction reflects memory maintenance and a trend toward mild improvement for cued words’ (two-tailed paired t-test, t(18)=1.72, p=0.1), whereas memory for uncued words significantly deteriorated after sleep (two-tailed paired t-test, t(18)=2.84, p=0.01). Nonparametric statistics further confirmed that memory was stable for cued words (Wilcoxon sign-rank=48.5, Z=-1.02, p=0.31) but significantly deteriorated for uncued words (Wilcoxon sign-rank=114, Z=-2.41, p=0.016). To further assess memory performance, we normalized post-sleep memory performance of each participant to her pre-sleep performance (set to 100%). The results (Fig. 2A) revealed that after sleep, memory for cued words (mean ± SEM=106% ± 17.5%, median=100%) was higher (t(18)=3.05, p=0.007; p=0.009 via Wilcoxon sign-rank test) than for uncued words (mean ± SEM=89.7% ± 17.0%, median=91.7%), and this was evident in most (13/19) participants. Bayes Factor for H1, was BF_10_ =4.391, further supporting the notion of improved memory for cued words.

**Figure 2.**
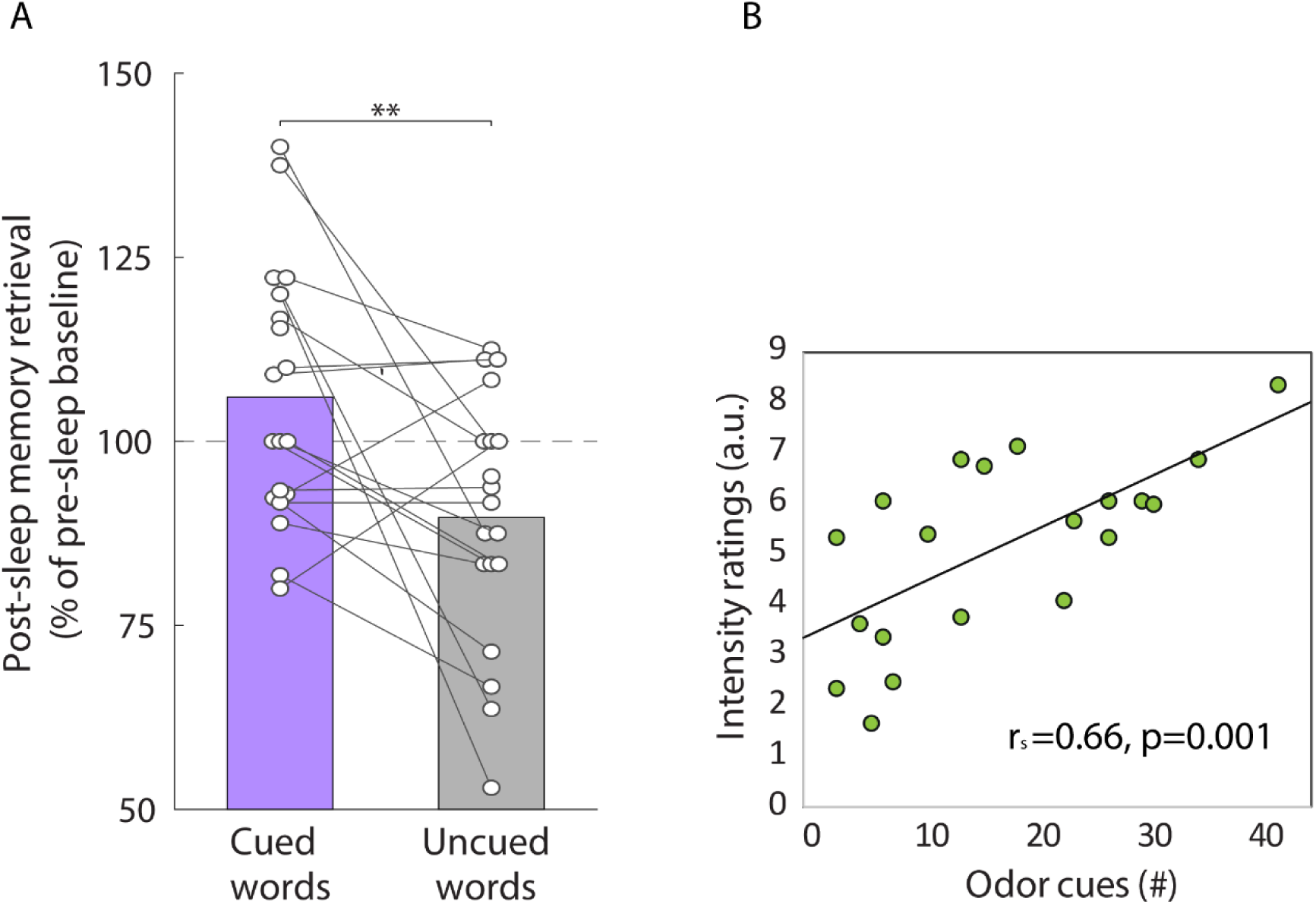
Behavioral effects of unilateral odor cueing during sleep. (A) Post-sleep memory retrieval (% of pre-sleep baseline) for cued word (purple, left) vs. uncued words (gray, right). Circles connected by lines mark data of individual participants. (B) Correlation between subjective odor intensity ratings following sleep (y-axis) as a function of the number of 30s odor stimulation epochs in N2 sleep.

To test whether odor presentation during sleep may influence additional cognitive aspects beyond excplicit memory, we measured the perceived intensity of the odor after sleep via subjective questionnaire ratings. Perceived odor intensity positively correlated with the number of 30s odor stimulation epochs in N2 sleep (r_s=_0.66, p=0.001, Fig. 2B). Memory or perceived intensity were not significantly different when comparing cueing in left vs. right nostrils, or words presented in left vs. right hemifields.

Together, the behavioral results indicate that unilateral odor cueing during sleep covertly affects the representation of odor cues, and improves memories for cued words associated with hemisphere-specific processing.

### Local TMR during sleep differentially modulates brain activity across hemispheres

To test how unilateral odor cueing modulated brain activity during sleep, we first evaluated the EEG spectral power changes during 30s odor stimulation epochs compared to baseline epochs, focusing on central electrodes (Fig. 3A & Methods). We found that odor stimulation significantly increased SW (<4Hz) and spindle (12-16Hz) power over both hemispheres (Fig. 3B: SW power in cued hemisphere: z=1.979, Wilcoxon sign-rank=158, p=0.048; SW power in uncued hemisphere: z=2.613, Wilcoxon sign-rank=175, p=0.009; spindle power in cued hemisphere: z=2.613, Wilcoxon sign-rank=175, p=0.009; spindle power in uncued hemisphere:, z=2.725, Wilcoxon sign-rank=169, p=0.017). A direct comparison between power changes across the two hemispheres revealed a significantly smaller increase (z=-2.725, Wilcoxon sign-rank=32, p=0.006) in SW power in the cued hemisphere compared to the uncued hemisphere (Fig. 3C). No significant difference was observed in spindle power elevation between hemispheres (p=0.65 via Wilcoxon sign-rank).

**Figure 3.**
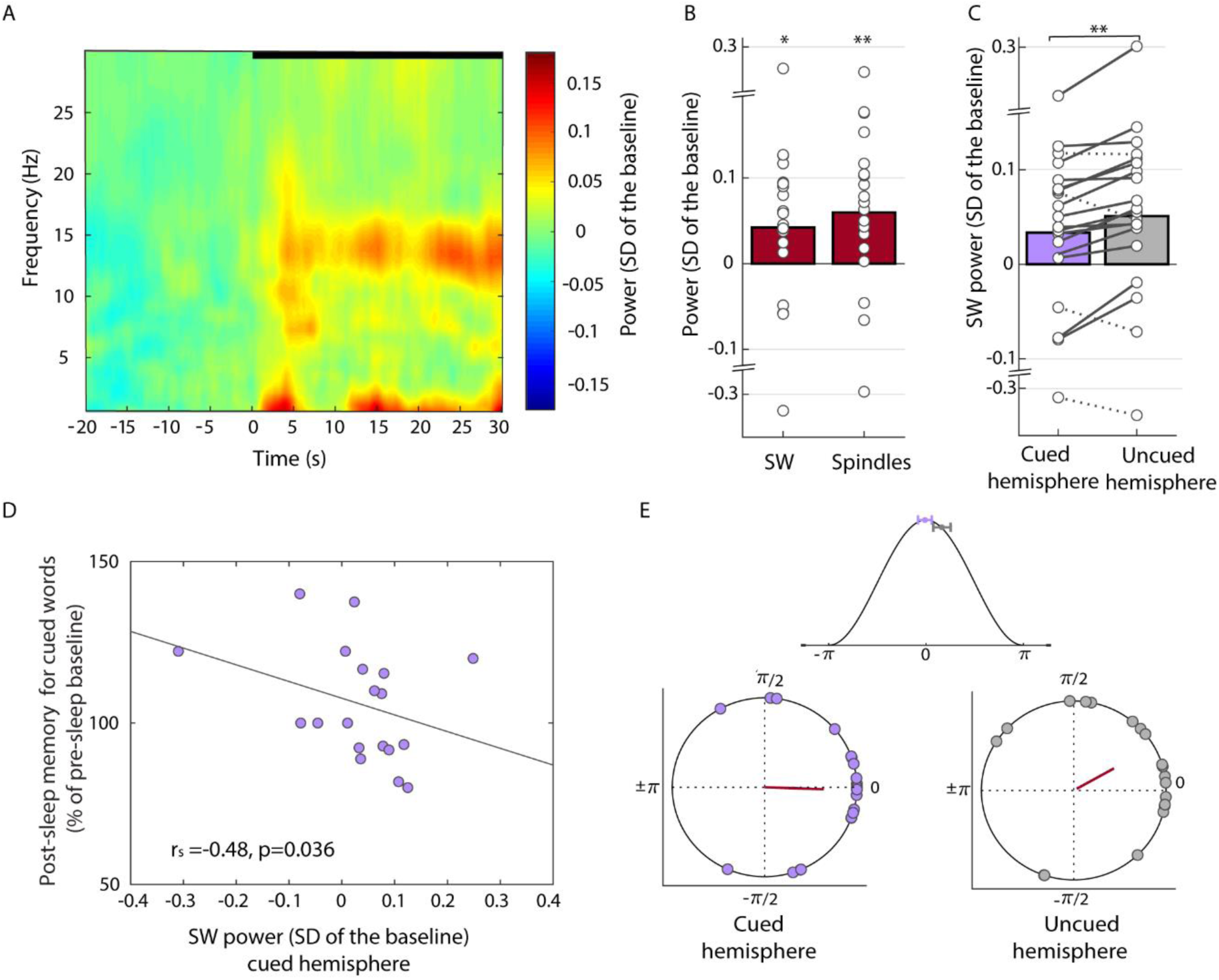
EEG effects of unilateral odor cueing during sleep. (A) Time-frequency decomposition of power changes in central EEG induced by odor cues during sleep (average of both hemispheres, n=1392 epochs, and 20 participants). Thick black bar on top represents 30s odor stimulation. Hot colors mark power increases in the SW (0.5-4Hz) and spindle (12-16Hz) frequency ranges (colorbar legend on right, in SD of baseline). (B) Power changes in central EEG for SW (left bar) and spindle (right bar) activities, averaged across both hemispheres. White circles mark data of individual participants. * p<0.05, ** p <0.01 compared to baseline. (C) Average change in SW power in the cued hemisphere (purple, left) vs. uncued hemisphere (gray, right) hemispheres. Circles connected by lines mark individual participant data. Solid/dashed lines mark lower/higher power in the cued hemisphere, respectively. ** p <0.01. (D) Scatter plot of post-sleep memory for cued words (as % of pre-sleep baseline, y-axis) vs. SW power in the cued hemisphere during odor cueing (as SD of baseline, x-axis) reveals significant negative correlation (r=-0.48, p=0.036). (E) Mean SW phase at time when sleep spindles occurred during odor cues (‘preferred phase’). Purple and gray circles denote preferred phase for cued hemisphere (left) and uncued hemisphere (right) for individual participants. Red bar marks the grand average phase (angle) and vector (radius). Top inset, grand average preferred phases and SEM (as horizontal bar) superimposed with schematic slow wave (cosine, peak=0°). Note earlier SW-spindle coupling around up-state peak in cued hemisphere.

We further found a relation between SWA changes and memory, whereby SWA in the cued hemisphere was negatively correlated with memory improvement for cued words (r_s_= −0.484, p=0.036, Fig. 2D). By contrast, odor-induced SWA in the uncued hemisphere did not correlate with memory improvement for uncued words (r_sp_=0.11, p=0.653). Analysis of EEG power changes in frontal electrodes yielded similar results (not shown). Thus, odor cues differentially modulate SWA in the cued vs. uncued hemispheres, and the smaller SWA increase in the cued hemisphere correlates with memory for select cued words processed in this hemisphere.

Finally, we tested whether unilateral odor cueing affected the temporal coupling between sleep spindles and slow wave up-states, implicated in memory consolidation [5, 43]. Analyzing the typical phase of slow waves at which sleep spindles preferentially occurred (Methods) we observed tight locking of spindles around SW peak (up-state): (cued hemisphere: resultant vector: 0.641, Rayleigh test for non-uniformity: z=7.35, p=3*10^-4^’; uncued hemisphere: resultant vector: 0.461, Rayleigh test for non-uniformity z=4.26, p=0.012). Additional nonparametric tests confirmed tight SW-spindle coupling, which was significantly different from a uniform phase distribution (p=0.0098 and p=0.03 for cued and uncued hemispheres, respectively, via Hodges-Ajne tests). A direct test comparing the mean SW phase at which sleep spindles occurred across the cued and uncued hemispheres during odor cues (Watson-Williams test for equal means, circular data t-test-equivalent, Methods) confirmed a significant difference in mean phase across hemispheres (F(1,37)=4.78; p=0.03). Additional nonparametric tests (similar to Kruskal-Wallis test for linear data) further confirmed the significance of this difference (p=0.003). Importantly, unilateral odor cueing differently modulated SW-spindle coupling across hemispheres (Fig 3E). While sleep spindles were time-locked to SW up-states over both hemispheres during odor stimulation, spindles occurred significantly earlier (closer to the positive peak of SWs) in the cued hemisphere (mean phase in cued hemisphere: =-1.971°, mean phase in uncued hemisphere=29.46°). Analysis of frontal EEG did not reveal significant differences in SW-spindle coupling as was found for central EEG (not shown). Together, we find that unilateral cueing led to earlier coupling between sleep spindles around slow wave up-states over central cortex and this occurred specifically over the cued hemisphere.

## Discussion

We used local TMR to enhance memories associated with processing in a single brain hemisphere. Odor presentation to a single nostril during sleep was intended to reestablish the context of learning before sleep, and led to differential effects of memory consolidation for select items. We observed improved memory for a subset of stimuli primarily processed in the cued hemisphere. In parallel, unilateral odor cues modulated regional sleep oscillations across the two hemispheres. Spindle activity and SWA increased in both hemispheres during odor stimulation, but SWA increase was lower in the cued hemisphere than in the uncued hemisphere. Furthermore, SWA increase in the cued hemisphere negatively correlated with memory improvement for the cued words. Temporal coupling between sleep spindles and slow wave up-states, implicated in sleep-dependent memory consolidation [1, 5], improved locally in the cued hemisphere compared to the uncued hemisphere. Together, the results demonstrate that local TMR in human sleep goes beyond global effects by selectively promoting consolidation of specific memories associated with regional sleep oscillations.

The memory effects observed here with local TMR are in line with TMR studies comparing contextual odor and vehicle cueing in sleep [10, 44]. In those studies, cued items show similar memory performance before and after sleep, whereas uncued items exhibit significant deterioration. Importantly, local TMR demonstrates such differences during the same sleep interval, thereby indicating that TMR acts locally on select engram representations. Post-sleep questionnaires verified there was no explicit awareness of odor cueing, as previously reported [10]. Although apparently not consciously perceived, the number of odor stimulation epochs during sleep significantly correlated with subsequent subjective ratings of odor intensity. This indicates that odor cueing during sleep covertly affects stimulus representation beyond its effects on memory consolidation. A previous study by Cox et al. (2014) [29] employed a between-subject experimental design using bilateral odor TMR. Two different odors were associated during learning with specific words presented in either left or right visual fields, and served as cues in separate sessions. They showed differential activity patterns across the two hemispheres during sleep, but did not find differential memory effects. Our local TMR enabled a within-session comparison that likely increased sensitivity and overcame variability across individuals and sleep sessions, revealing significant differences in memory for cued vs. uncued items.

Another approach to modulate consolidation of specific human memories in a local manner is to perform closed-loop stimulation during sleep that is time-locked with regional oscillations. Such stimulation increases spindle activity locally and enhances motor memory consolidation [45], and can disrupt SWA locally and interfere with motor learning [46]. These studies compared brain activity across two different sessions, where considerable variability exists in sleep, brain activity, and memory processing. Our method goes beyond variability in global sleep factors, by modulating local oscillations and improving memory consolidation of corresponding items selectively.

The local TMR effects on cortical sleep oscillations extend previous studies showing that olfactory stimulation during sleep enhances SWA [23, 44]. We found that SWA increase was lower in the cued hemisphere and negatively correlates with memory for cued items (but not for uncued items). The processes mediating smaller elevation in SWA in the cued hemisphere remain unclear, but one possible explanation could be that unilateral odor stimulation elicited selective processing in the cued hemisphere associated with EEG desynchronization and lower SWA. In other words, activity in the cued hemisphere may represent a combination of global SWA increases and local processing that reduces synchronized slow activities. The fact that a lower SWA increase correlates with better memory for cued stimuli may reflect a “penetration” of the odor stimulus that efficiently modulates ongoing activities in the cued hemisphere.

The mechanisms mediating the memory benefits of TMR are not fully established, but may involve sensory-triggered hippocampal reactivation and hippocampal-neocortical dialogue [1, 4], as recently reported in rodent studies [47]. Our results support the notion that temporal coupling between sleep spindles and slow wave up-states is a non-invasive proxy for effective sleep-dependent memory consolidation [1, 5, 43, 48–51]. Going beyond correlative evidence, we find that local TMR causally elicits tighter SW-spindle coupling in the cued hemisphere, highlighting the regional quality of sleep oscillations and their relation to memory consolidation [27, 29, 30, 33, 52, 53]. We find tighter SW-spindle coupling in central (but not in frontal) electrodes of the cued hemisphere, in line with findings that posterior spindles are preferentially involved in memory consolidation [29, 43, 51].

In summary, we report here a novel non-invasive technique for local TMR of specific regional memories, opening new avenues for sleep and memory research, as well as potential clinical applications. For example, unilateral sleep TMR could be used to modulate different components of traumatic memories in post-traumatic stress disorder (PTSD) that are lateralized [54] or assist rehabilitation in individuals with lateralized brain damage such as unilateral stroke.

## Methods

### Participants

Thirty two healthy participants took part in this study (18 women, mean age 27.87 ± 3.8 years). All participants reported no history of psychiatric, neurological, sleep or respiratory disorders. All were good habitual sleepers and native Hebrew speakers. The participants provided written informed consent to the procedures approved by the committee for protection of human participants at the Sleep Disorders Laboratory, Loewenstein Rehabilitation Hospital, Raanana. Participants received monetary compensation for their participation.

On the day of the nap experiment, participants were instructed to wake up no later than 8.30 AM and to eat their lunch before the experimental session began in the early afternoon. They were further instructed to limit their caffeinated beverages to one, and to avoid ingesting alcohol during the experimental day. Exclusion criteria were insufficient (<20 minute) time in NREM sleep (10 participants were excluded), or technical problems either in the odor delivery system (2 participants excluded) or with behavioral memory data (one participant excluded due to technical failure). Subsequent analysis was performed on all behavioral data (n=19, 11 women, mean±SD age=27.4 ± 2.9 years, range 21-30) and on all sleep EEG data (20 participants, 11 women, mean±SD=27.4 ± 2.8 years, range 21-30).

### Experimental procedure

An overview of the experimental procedure is presented in Fig 1A,B. Participants arrived at the lab in the early afternoon (between 12:00 and 14:00). The experimental room was coated with stainless steel to prevent ambient odor adhesion. In addition, a high-efficiency air filter further assured an odor-free environment. Next, the participants performed the training session of the spatial memory task while bilateral odor stimulation was delivered through a nasal mask covering both nostrils. Training was followed ~5 minutes later by a memory retrieval test without an odor mask. Next, after fitting of the polysomnographic electrodes and a separate nostril mask, participants were left in bed undisturbed to sleep for two hours (Fig. 1E). Throughout the nap, participants were monitored online from the adjacent control room through a one-sided window, and polysomnographic measures were inspected online at all times (Supplementary Fig. S1). During NREM sleep, a rose odor was presented to one nostril (left vs. right nostril counterbalanced across participants, n=11 vs. 9, respectively). Upon awakening and a short recovery (~15 min), participants performed the post-sleep memory retrieval test. This test was followed by intensity rating of the same odor used in the task and sleep. Total duration of the experimental session was approximately 4.5 hours.

### Memory task

The memory task, as well as the retrieval test and the distraction task were implemented in <u>MatLab</u> using <u>Psychophysics toolbox</u> extensions [55–57]. The memory task was adapted from Cox et al. (2014) [29] with minor changes. In each trial, a different word appeared in the center of the screen, and then moved toward one of eight targets arranged in circle on the screen. The participants were instructed to memorize the associations between the words and the targets (Fig. 1B). Each target was associated to the same number of words. Word stimuli consisted of common Hebrew concrete nouns (3-5 letters) obtained from stimuli in [58] and their frequency was controlled for in <u>The Word-Frequency Database For Printed Hebrew</u>[59]. Thirteen participants memorized the word-target associations of 32 words, and six participants memorized 48 associations. All results were derived after collapsing these two groups together, reporting the percent of remembered associations out of all learned associations. The targets were arranged in a circle on the screen, with four targets located in the right visual field and four targets in the left visual field, to elicit processing in the left and right hemispheres, respectively (Fig. 1D). Participants were instructed to fixate on a white fixation cross in the center of the screen at all times. The circular targets arrangement was used to render an implicit left/right division of the targets that was transparent to the participants (Fig. 1B). Words appeared in two blocks and repeated in six learning rounds. The number of learning rounds was determined following a pilot experiment to form a strong, yet imperfect memory of the words, aiming for success rate around 60% before sleep (chance level is 12.5%). In addition, a pilot experiment with eye tracking <u>(Eyelink-1000, SR Research</u>) verified the ability of participants to focus their eyes on the fixation cross, and not let their gaze follow the moving word. (Supplementary Fig. S1). A rose scent was delivered to both nostrils via a nasal mask throughout the encoding phase (Fig. 1B). Memory was first tested immediately following the training session (pre-sleep memory test), and again later within ~15 min of the end of the nap (post-sleep memory test). During these tests, all targets appeared on the screen and words were shown sequentially in a random order. The participants had 10s to select a target and instructed to guess the correct target if cannot remember. After 10s, if none of the targets were selected, the next trial began (Fig. 1C). No feedback was provided regarding accuracy. Odors were not delivered during the memory test. Supplementary material provides additional details regarding the memory task.

### Olfactory stimulation

A computer controlled air-dilution olfactometer [60, 61] was located in an adjacent room, with tubes entering the experimental room through waveguides to prevent potential non-olfactory signals from the olfactometer body. These tubes ended in a small nasal mask (different masks for wake and sleep, as described below). The olfactometer delivered a constant flow of clean air, at a rate of 5 liters per minute (lpm). The airflow to the nose was constantly vacuumed away at the same rate, preventing accumulation or lingering of odor. Odorant pulses were embedded in the flow at certain times of the experiment. High temporal resolution of the stimulus was achieved using a railroad manifold that enabled switching between odor and clean air close to the nasal mask as described in [60]. The odorant (Phenyl-ethyl alcohol, CAS Number 60-12-8 Sigma-Aldrich, generally perceived as “rose”) was chosen because it has been shown to minimally activate trigeminal pathways [62]. This odorant has been used successfully as a stimulus in a number of sleep experiments [10, 63]. Task mask: The mask (Respironics, contour deluxe SU) covered both nostrils. Odor and clean air stimulation enter the mask from the olfactometer via a Teflon tube. A vacuum tube line evacuated the air, and two small probes were attached to spirometer for measuring respiration. Nasal sleep mask: The sleep mask was designed and built in-house especially for this experiment. Its uniqueness is in its complete separation of the olfactory environment between nostrils. The nasal part comprised of a silicone nasal pillow (ResMed, Swift™ FX Nasal Pillow CPAP Mask). Two stainless steel tubes embedded in Teflon adapters were inserted to holes drilled in the nasal pillow to separate the airflows to each nostril. These inlets are connected to wide plastic pipelines from a BIODEX Disposable Xenon-133 Rebreathing Systems. These pipelines open to room air on the other side and odor air and vacuum tubes within them control the composition of air inside. The participant breathes passively the air in the void within these pipelines. Air tube in one pipeline delivered constant flow of clean air while the air tube on the other pipeline delivered clean air/odor stimulation according to the operator's instructions. Vacuum lines evacuated the air in the pipelines. Two spirometer probes, one in each pipeline, measures respiration separately from each nostril. Tubes coming out of the mask were suspended from a rail close to the celling to prevent discomfort in the sleeping participant due to weight of gear (Fig. 1E, supplementary Fig. S3).

### Odor cueing in sleep

Through all sleep stages, the experimenter monitored the polysomnographic measures to identify the sleep stages and deliver odor stimulation accordingly. Odor stimulation started after at least 10 minutes of sleep and for as long as the participant remained in NREM sleep. The stimulation was stopped upon any polysomnographic signs of arousal, awakening, or REM sleep. Stimulation followed an alternating 30s-on/30s-off pattern (referred to as ‘odor on’ and ‘odor off’ periods throughout) to reduce habituation. Alternating between these phases was done without any visual, auditory, temperature or other non-olfactory hints that could be perceived by participants.

### Polysomnography and sleep scoring

Physiological measurements were acquired during sleep with a PowerLab Monitoring System (ML880 ADInstruments, Bella Vista, NSW, Australia) at a sampling rate of 1kHz with a 50Hz notch filter. Electroencephalogram (EEG) was obtained with four circular electrodes located at positions C3, C4, F3 and F4 according to the 10-20 system, referenced to electrodes on the opposite mastoids: A2 and A1. Electroocculogram (EOG) was obtained with two circular Ag/AgCl conductive adhesive electrodes, placed 1 cm above or below and laterally of each eye and referenced to electrodes on the opposite mastoids. Electromyogram (EMG) was obtained with two circular Ag/AgCl conductive adhesive electrodes, located bilaterally adjacent to the submentalis muscles. EEG, EOG and EMG signals were amplified with a preamplifier (Octal Bio Amp ML138, ADInstruments). Electrocardiogram was obtained with two circular Ag/AgCl conductive adhesive electrodes, placed on left and right sides of the abdomen and referenced to a ground electrode placed on the left foot. Nasal respiration was measured separately from each nostril using two spirometers (FE141, AdInstruments). A representative example of polysomnography data is presented in Supplementary Fig. S1. Sleep stages were visually scored off-line by two independent scorers according to the 2012 updated American Academy of Sleep Medicine (AASM) Manual [64]. Sleep parameters, including sleep latency, total sleep time and time spent in different sleep stages, were calculated (Supplementary Table ST1).

### EEG data analysis

For all the EEG analyses, custom Matlab scripts were combined with functions from EEGLAB toolbox [65] To remove noise, the EEG signal was first filtered with a FIR high-pass filter at 0.5Hz using a Hamming window and transition band of 0.5Hz. The signal was then filtered with FIR low-pass filter at 30Hz to remove high-frequency noise with a Hamming window and transition band of 2Hz. After filtering, the signal was down-sampled to 250Hz and divided to REM and NREM epochs. Noisy channels (one channel from one individual) and epochs (18/11/2 epochs from 3 individuals) were removed from analysis. All the ensuing EEG analyses were conducted on NREM epochs after these pre-processing steps.

Dynamic spectral analysis using event-related spectral perturbation (ERSP) was implemented as follows: For each channel separately, we extracted 55s epochs around the odor cues (25s ‘odor-off’ baseline period followed by 30s ‘odor-on’ period, avoiding the few seconds following odor termination to allow activity to return to baseline levels). The signal was divided to 0.5Hz frequency bins (59 bins 0.5-30Hz) using FIR filter and Hamming window. For each frequency bin separately, we performed a Hilbert transformation and then normalized the preceding 25s baseline:

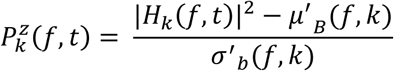

Where *H*_*k*_(*f*,*t*) is the Hilbert tranfromation for each time point *t* in frequency bin *f*, and trial *k* (odor cue). *μ*^′^_*B*_(*f*, *k*) is the mean baseline (‘odor–off’ period) spectral estimates for trial (odor cue) *k* and frequency bin *f*, and *σ*^′^_b_(*f*, *k*) is the spectral estimate SD for the baseline period of trial *k* at frequency bin *f* [66] Then, for each participant separately, we took the median normalized power across all odor cues. For visualization, averaged ERSP across all participants and channels are shown in Fig. 2A. To further quantify the modulation in power as a result of odor cues, and to compare this modulation between the hemispheres, we evaluated the power in the two frequency bands of interest: SW range (0.5-4Hz), and spindle range (12-16Hz).

#### Detection of sleep spindle events

For detection of sleep spindles and phase amplitude coupling (PAC) analysis with SW, we used unfiltered raw data to avoid potential phase distortion that could affect PAC analysis. Other preprocessing steps were identical. Discrete spindles were detected automatically in artifact-free NREM data using a Matlab custom script based on methods described previously [29, 33, 44], with minor adaptations. First, the NREM EEG signal was bandpass-filtered in the spindle range (12-16Hz), using a zero-phase two way least-squares FIR filter with transition band width of 2Hz. The signal envelope was calculated with a Hilbert transform. The amplitude was further smoothed using a Gaussian kernel (σ=40ms), to avoid multiple crossings of thresholds within the same spindle event. Events were detected when the envelope amplitude crossed the detection threshold of mean + 3SD, computed for each electrode and each participant separately, across all the NREM sleep signal. Multiple events detected within 1s were merged. The event start/end times were determined when the signal crossed the threshold mean + 1 SD of spindle power across NREM sleep. Events with duration between 0.5s and 2s were considered for further analysis (Supplementary Fig. S4). To further reduce the probability of detecting spurious spindles, spectral power of the detected events was calculated using a Fast Fourier Transform (FFT) on the noise-free raw signal. Only events with peak spectral-power within the spindle-frequency range (12-16Hz) compared to adjacent frequency bands (7-9Hz and 18-20Hz) were considered, to avoid events with broadband power increases.

#### SW-spindle phase amplitude coupling

For detecting the preferred SW phase at which sleep spindles occurred, we used the subset of spindles detected (above) that occurred during unilateral odor cueing (‘odor on’ periods), and used the maximal amplitude of spindle envelope as the spindle occurrence timing. SW phase corresponding to the spindle peak amplitude was computed by first filtering the raw signal in the (0.5-2Hz) frequency band using a zero-phase two-way least-squares FIR filter with low transition bandwidth of 0.08Hz and high transition bandwidth of 0.3Hz, and then extracted the instantaneous phase using the Hilbert Transformation. For each participant separately, we then calculated the ‘preferred phase’ as a circular mean of all SW phases for corresponding to sleep spindle events.

### Statistical analysis

Statistical analyses were performed within-subjects using custom Matlab scripts. For behavioral data and for phase-amplitude coupling, both parametric and nonparametric statistics were used. Change in memory performance over sleep was assessed with two-way repeated measures ANOVA using time (pre-sleep/post-sleep) and cueing (cued/uncued) as factors, followed by post hoc comparisons using two-tailed paired t-test, as well as Wilcoxon sign-rank tests. Next, individual post-sleep memory performance was normalized to pre-sleep performance (pre-sleep performance was set to 100%), and change in memory for words was compared between sides with a paired t-test as well as Wilcoxon sign-rank test. We used Spearman correlations to assess the correlation between the number of odor stimulation epochs and perceived odor intensity.

For EEG power spectrum analysis, non-parametric statistical tests were used. To test the effect of odor cueing on EEG power we performed one-sample Wilcoxon sign-rank tests on normalized EEG activity during odor cueing. As EEG power was normalized to the ‘odor-off’ baseline in each cueing epoch separately, values different than zero indicate significant odor-induced modulation. Next, we compared the induced EEG power between hemispheres (cued and uncued) separately for SW (0.5-4Hz) and spindle (12-16Hz) frequency bands, using Wilcoxon tests. We used Spearman correlations to assess the correlation between SW power and memory improvement.

In SW-spindle coupling analyses, circular statistics were calculated using the CircStat toolbox [67]. The distribution of ‘preferred phases’ across participants was tested against uniformity with both parametric (Rayleigh) and nonparametric (Hodges-Ajne) tests. To statistically compare the odor-induced ‘preferred phases’ in a pairwise manner across cued and uncued hemisphere data while minimizing inter-subject variability, we normalized the ‘preferred phase’ in each participant separately by subtracting the average phase of both hemispheres from both the cued and from the uncued hemispheres preferred phases. We then tested for equal circular means using Watson-Williams test (circular t-test equivalent) and with non-parametric circular Kruskal-Wallis equivalent.

## Acknowledgements

We thank Aharon Weissbrod for his help in designing and building the nasal masks and olfactory stimulation setup. Prof. Rony Paz for his support. Dr. Motti Ratmansky from Loewenstein Rehabilitation Hospital. Members of Sobel/Nir/Paz labs and Dudai lab alumni for discussions and suggestions. This work was supported by the Israel Science Foundation (ISF) grant 51/11 (I-CORE cognitive sciences, YN).

## Author contributions

E.B. conceived of research and designed experiments with supervision from Y.D. and N.S. E.B., E.L. and N.S. designed and built the experimental setup

E.B. collected data.

E.B. analyzed data with the help of A.A., O.P. and supervision from Y.N.

E.B. and Y.N. wrote the manuscript.

All authors provided ongoing critical review of results and commented on the manuscript.

## Supplementary Material and Figures

### Training Session

The learning and memory task was adapted from Cox et al. (2014) [29] with minor changes. Each training block started with a 1600ms fixation period (1400ms for participants that learnt 48 words) with a white cross in the middle of the screen and all eight targets appearing as small gray octagons with green borders (Fig. 1B). Then, each trial started with a fixation period of either 200ms, 400ms, or 600ms (pseudorandom). Next, a word appeared at the center of the screen for 800ms, followed by transient enlargement of one target location to medium size for 400ms. Next, the word moved toward that target while its font size decreased and the target octagon increased further to a large size for 150ms. Finally, the word appeared clearly inside the target octagon for another 400ms before the next trial started. The words were presented serially in two blocks (half of the words in each block). The blocks were biased to either the left or right visual hemifield, such that in one block 75% of the words moved to left targets and in the other block 75% of the words moved to right targets. Overall, each target was associated with the same number of words. The word order, the word-target associations, and the word-block assignments were randomized across participants, with one restriction - that the same target location could not appear twice consecutively, to limit the formation of cognitive links between words associated with the same location. Participants were instructed to maintain fixation at the center of the screen at all times. Presentation of a full word block lasted 36s. Each block was followed by a non-hippocampal “distraction” task (counting back out loud in steps of three starting from a three digit number appearing on the screen, as in Cox et al. (2014) [29] to minimize active memorizing and preserve the memory to be as visual as possible. Odor stimulation occurred during the word blocks but not during the distraction tasks or during breaks between rounds. After each block, participants reported whether they could smell the odor during the task or not, by pressing on one out of two keyboard buttons. Two blocks (consisting of all of the learned words) together with corresponding distraction tasks made up one learning round. Block order (left vs. right hemifield bias) in each learning round was also randomized. Each participant completed six learning rounds separated by one minute breaks. The total duration of the learning task encoding phase was about 30 minutes.

### Memory Retrieval Test

Each test trial (Fig. 1C) started with presentation of all eight targets together with a white central fixation cross for either 200ms, 400ms, or 600ms (pseudorandom). Then, a word was presented in the middle of the screen for 2500ms. Next, a white point curser replaced the word and the participants could move it using the computer touchpad toward the target they thought was associated with this word in the encoding phase. When the cursor was close to a target, the octagon was enlarged to medium size, and after target selection with the touchpad button press, the octagon grew further to large size and remained that way for 2500ms until the next trial started. The total duration of the retrieval test was about 10 min.

### Pilot experiment and eye tracking

Before running the main experiment, a pilot experiment (n=28 participants) was performed to assess (i) how many learning rounds were needed to set the mean success rate around 60%, and (ii) the ability of the participants to fixate successfully without gaze following the moving word. We recorded eye movements at 500Hz using an infrared Eyelink-1000 eye tracking system (SR-Research), at a sampling rate of 500 Hz. Figure S1 shows eye movement heat map histograms of 28 individual participants demonstrating successful fixation.

---

**Figure S1.**
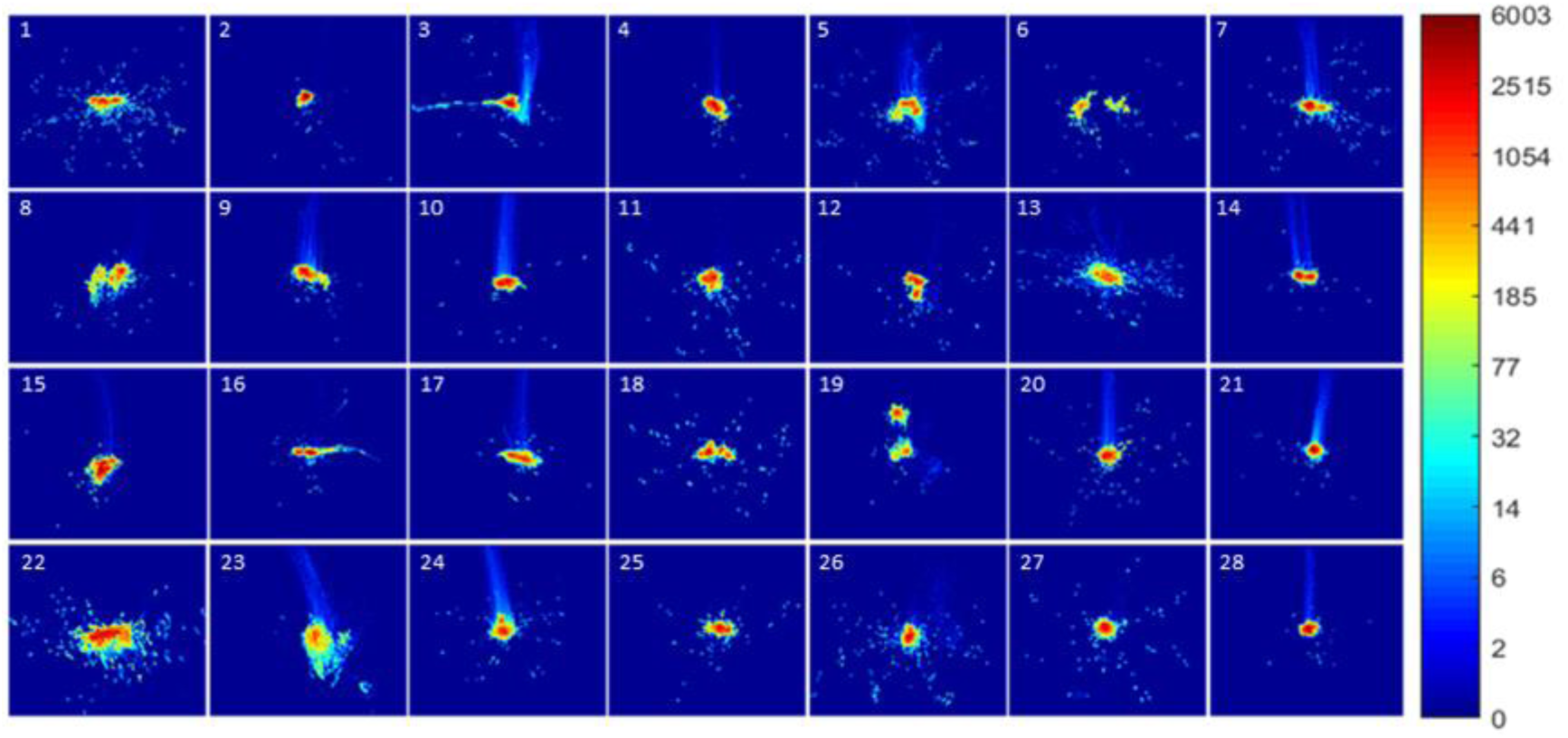
Eye tracking. Eye movement heat map histograms of individual participants (n=28). Colors (colorbar legend on right) represent the number of fixations the participant made at every bin (250×250 screen pixels). Participants 6, 8, 12, 19, 20 showing two noncontiguous hotspots represent a camera position change in the middle of the experiment due to a change in the position of the eye after the brake between the learning rounds.

**Figure S2.**
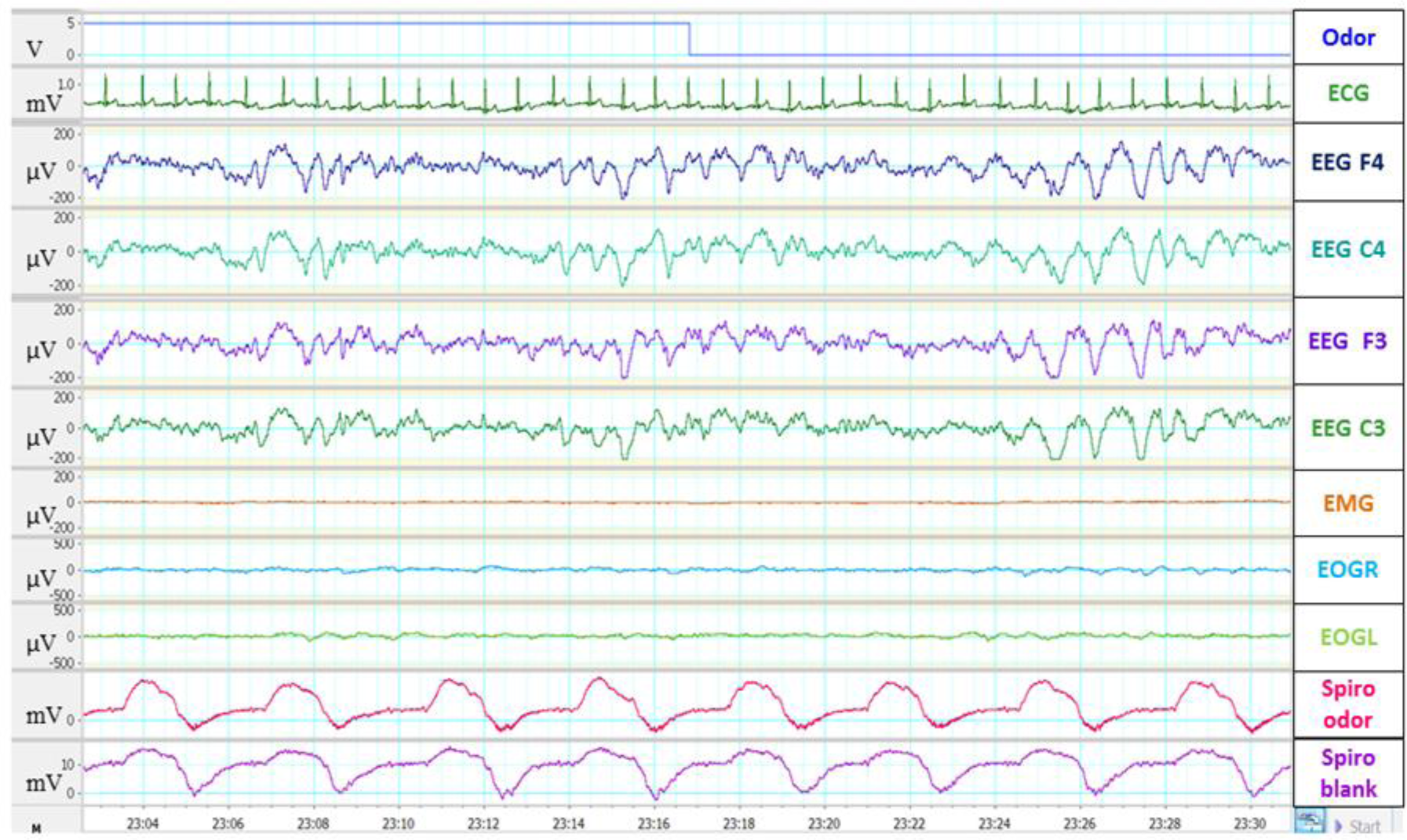
Example of 30s polysomnography recording from single participant during NREM sleep. Rows (top to bottom) show odor stimulus (blue; up phase - on, down phase – off), ECG (green), right frontal F4 EEG (dark blue), right central C4 EEG (cyan), left frontal F3 EEG (purple), left central C3 EEG (green), EMG (orange), right EOG (light blue), left EOG (light green), respiration in the stimulated nostril (pink), and respiration in the unstimulated nostril (purple).

**Figure S3.**
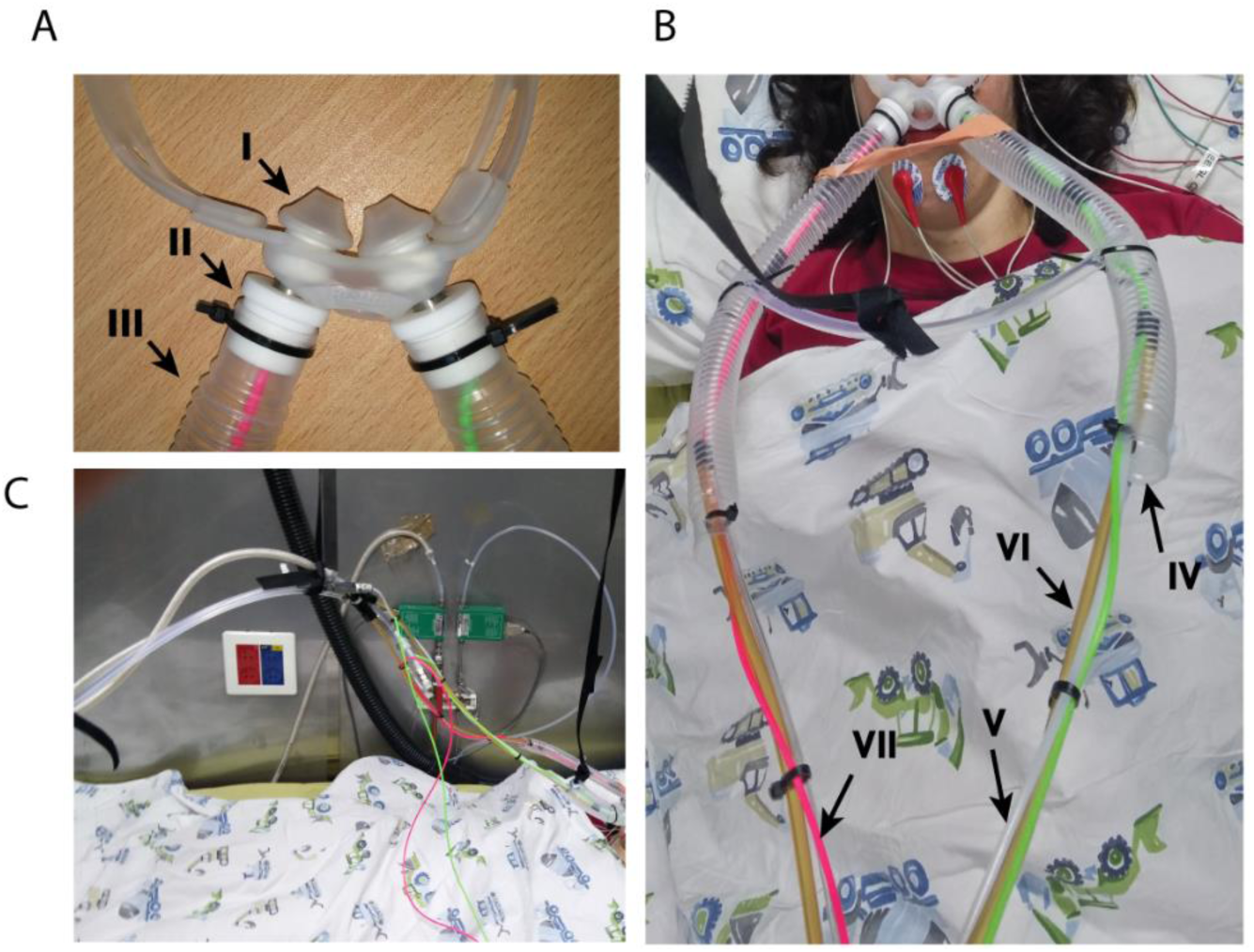
The nasal sleep mask enabling unilateral odor stimulation. (A) The nasal part: I. Nasal pillow from Swift™ FX Nasal Pillow CPAP Mask. II. Stainless still and Teflon inlets. III. Plastic pipeline from Disposable Xenon Rebreathing Systems Biodex. (B) The tubes construction: IV. The second end of the pipelines is open to room air. V. Clean air\odor line. VI. Vacuum line. VII. Spirometer probe (green and pink Tygon cannulas). (C) Tubes coming out of the mask were hanged from a ceiling rail to prevent discomfort on the sleeping participant due to weight of gear.

**Figure S4.**
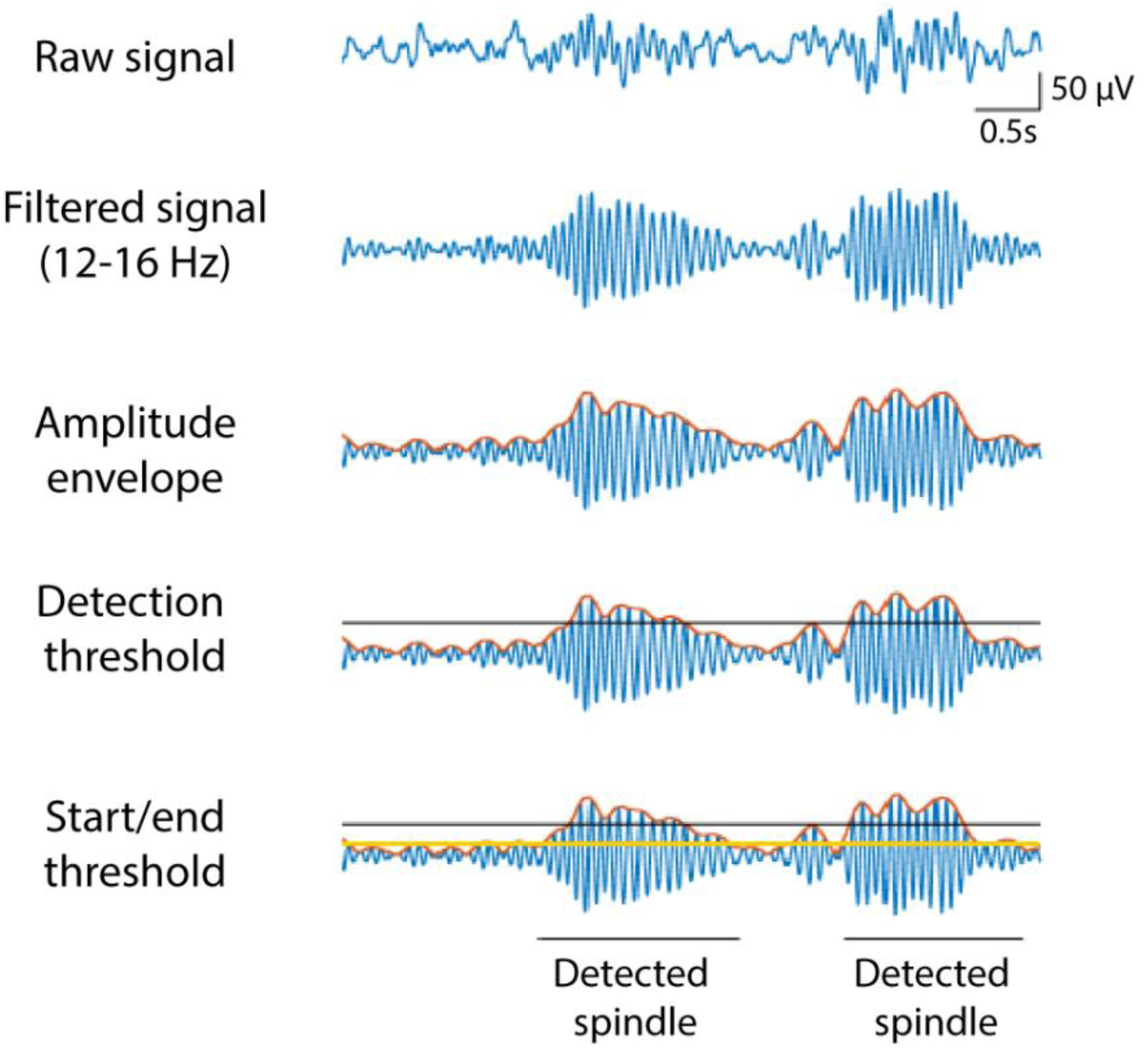
Detection of sleep spindles Illustration of spindle detection process. Raw EEG (top row) was filtered in the spindle frequency range (12-16Hz, second row). The amplitude envelope is extracted using Hilbert transform (red trace, third row). A spindle is detected whenever the envelope crossed a detection threshold of mean + 3SD of spindle power across NREM sleep (horizontal black line, fourth row). A start/end threshold is set at mean + 1SD of spindle power across NREM sleep (horizontal orange line, fifth row).

**Table ST1.**
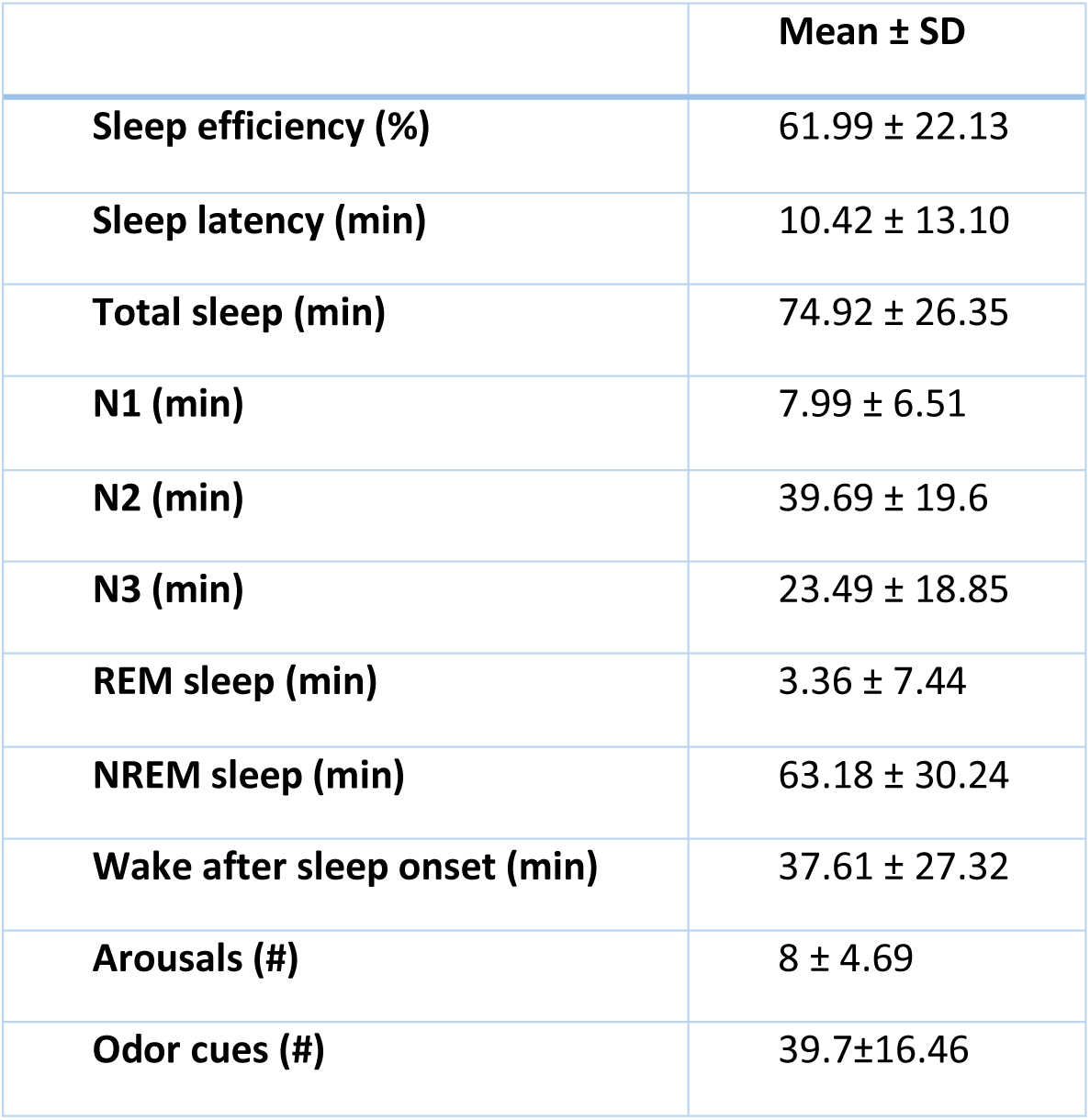
Sleep architecture parametersM. Sleep architecture parameters (mean ± SD), and the average number of odor cues for all participants included in the analysis (N=20).

|  | **Mean ± SD** |
| --- | --- |
| **Sleep efficiency (%)** | 61.99 ± 22.13 |
| **Sleep latency (min)** | 10.42 ± 13.10 |
| **Total sleep (min)** | 74.92 ± 26.35 |
| **N1 (min)** | 7.99 ± 6.51 |
| **N2 (min)** | 39.69 ± 19.6 |
| **N3 (min)** | 23.49 ± 18.85 |
| **REM sleep (min)** | 3.36 ± 7.44 |
| **NREM sleep (min)** | 63.18 ± 30.24 |
| **Wake after sleep onset (min)** | 37.61 ± 27.32 |
| **Arousals (#)** | 8 ± 4.69 |
| **Odor cues (#)** | 39.7±16.46 |

